# Motor adaptation does not differ when a perturbation is introduced abruptly or gradually

**DOI:** 10.1101/2022.12.22.521654

**Authors:** Ambika Bansal, Bernard Marius ’t Hart, Udai Chauhan, Thomas Eggert, Andreas Straube, Denise YP Henriques

## Abstract

People continuously adapt their movements to ever-changing circumstances, and particularly in skills training and rehabilitation, it is crucial that we understand how to optimize implicit adaptation in order for these processes to require as little conscious effort as possible. Although it is generally assumed that the way to do this is by introducing perturbations gradually, the literature is ambivalent on the effectiveness of this approach. Here we test whether there are differences in motor performance when adapting to an abrupt compared to a ramped visuomotor rotation. Using a within-subjects design, we test this question under 3 different rotation sizes: 30°, 45°, and 60°, as well as in 3 different populations: younger adults, older adults, and patients with mild cerebellar ataxia. We find no significant differences in either the behavioural outcomes, or model fits, between abrupt and gradual learning across any of the different conditions. Neither age, nor cerebellar ataxia had any significant effect on motor adaptation either. These findings together indicate that motor adaptation is not modulated by introducing a perturbation abruptly compared to gradually, and is also unaffected by age or mild cerebellar ataxia.

## INTRODUCTION

One of the most fundamental functions of the human brain is to adapt movements to our everchanging environment or body. Understanding how to reduce the effort it takes to make these adaptations, by capitalizing on our powerful implicit learning processes, will be highly beneficial to rehabilitation and skills training. Here we ask if adapting to small but gradual changes compared to abrupt changes modulates the effort it takes to adapt to them. We look at perturbations of different sizes and the effects of age and mild cerebellar ataxia on implicit adaptation.

### Abrupt versus Gradual Motor Learning

It is assumed that adapting to an abrupt perturbation is more effortful and explicit, whereas a gradually introduced perturbation may be more implicit (Taylor et al., 2014; Bond & Taylor, 2015). Gradually and abruptly introduced perturbations do sometimes result in different aftereffects, a hallmark of implicit learning, in visuomotor adaptation (Kagerer et al., 1997; Ingram et al., 2000), force-field adaptation (Kluzik et al., 2008), and prism adaptation (Michel et al., 2007). However, these effects do not generally replicate (Buch et al., 2003, Klassen et al., 2005), even when done by the same lab (Kagerer et al., 2006). Three recent papers have tested the notion that the way a perturbation is introduced affects the underlying learning processes differently in younger adults (Coltman et al., 2021, Alhussein et al., 2019), and in cerebellar patients (Hulst et al., 2021), and here we probe these questions further.

### Two-Rate Model of Motor Learning

A common model used to describe motor adaptation, and explain the rebound phenomena, consists of two processes, a fast and slow process (Smith et al., 2006). The fast process is quick to learn, but also quick to forget, whereas the slow process learns much slower, but also retains the learnt adaptation for much longer. The rebound phenomena, which proves that there is some retention of an initial perturbation that persists even after adapting to a second perturbation, may suggest the contribution of multiple learning mechanisms (Smith et al., 2006). The fast and slow process of the two-rate model have been suggested to map onto the explicit and implicit processes of learning, respectively (McDougle et al., 2015). If there is a greater contribution of implicit learning when a perturbation is introduced gradually, then we should see a greater contribution of the slow process as well.

### Motor Learning in Older Adults and People with Cerebellar Ataxia

The cerebellum plays a crucial role in our implicit motor learning (Hull, 2020). Although the functional deficits for people with cerebellar damage are still unclear, people with cerebellar ataxia can use an explicit strategy to compensate in making adaptive movements (Taylor et al., 2010). Previous research has found greater aftereffects when using a gradual perturbation schedule in people with severe cerebellar ataxia (Criscimagna-Hemminger et al. 2010). However, these findings were not always replicated in later studies (Gibo et al., 2013, Schlerf et al., 2013). Other work has found that aging has an analogous pattern of degeneration to that of people with cerebellar degenerative disease, and in some cases has been used as a model system of cerebellar disease (Hulst et al., 2015). Although there is evidence to show that the cognitive processes that decline with aging affect explicit learning processes, this deficit can be compensated with implict learning (Vandevoorde et al., 2019, Vachon et al., 2020).

### Objectives

The main objective of this study was to examine any differences in motor performance when responding to a perturbation that is introduced abruptly compared to ramped. We originally hypothesized that these different perturbation schedules would result in different levels of adaptation, and thus two-rate model fits. This hypothesis was based on two assumptions: (1) the fast and slow processes of the two-rate model map onto explicit and implicit motor learning, respectively, and (2) abruptly and gradually introduced perturbations elicit different amounts of explicit and implicit learning. We also predicted that performance and model fits could be further modulated by age, which may increase implicit adaptation, and by mild cerebellar ataxia, which may increase explicit adaptation.

## METHODS

## Experiment 1: Rotation Size

### Participants

Sixty-four students from York University participated in this experiment. There were thirty subjects who participated in the ‘younger 30°’ group (5 lefthanded, 8 males), and twenty-nine participants who participated in the ‘younger 60°’ (1 lefthanded, 9 males). Of this, 7 participants were removed from the 30° group, and 5 participants were removed from the 60° group, as they did not adapt to at least one of the perturbations. The 30° group consisted of subjects ages 20 +/-4 years old, and the 60° group were ages 20 +/-2 years old. All participants reported having normal or corrected-to-normal vision. The protocols used in this study were approved by the York Human Participants Review Sub-committee and by the Ethics Committee of the Ludwig-Maximilians University, Munich (559-15). All participants gave prior informed written consent, and were naive to the purpose of the study. Participants were recruited using the York University undergraduate research pools, and were given course credit for participation.

#### Apparatus

The equipment used in this experiment was a laptop (Dell Inc.), computer monitor (Dell Inc. 20”, 30 Hz refresh rate, 1680 × 1050 resolution), mirror, tablet (Wacom Intuos Pro, 311 × 216 mm), and stylus. The visual stimuli were projected from the downward facing monitor onto the mirror, such that the stimuli were perceived to be in the same horizontal plane as the tablet below (Figure 1).

**Figure 1.**
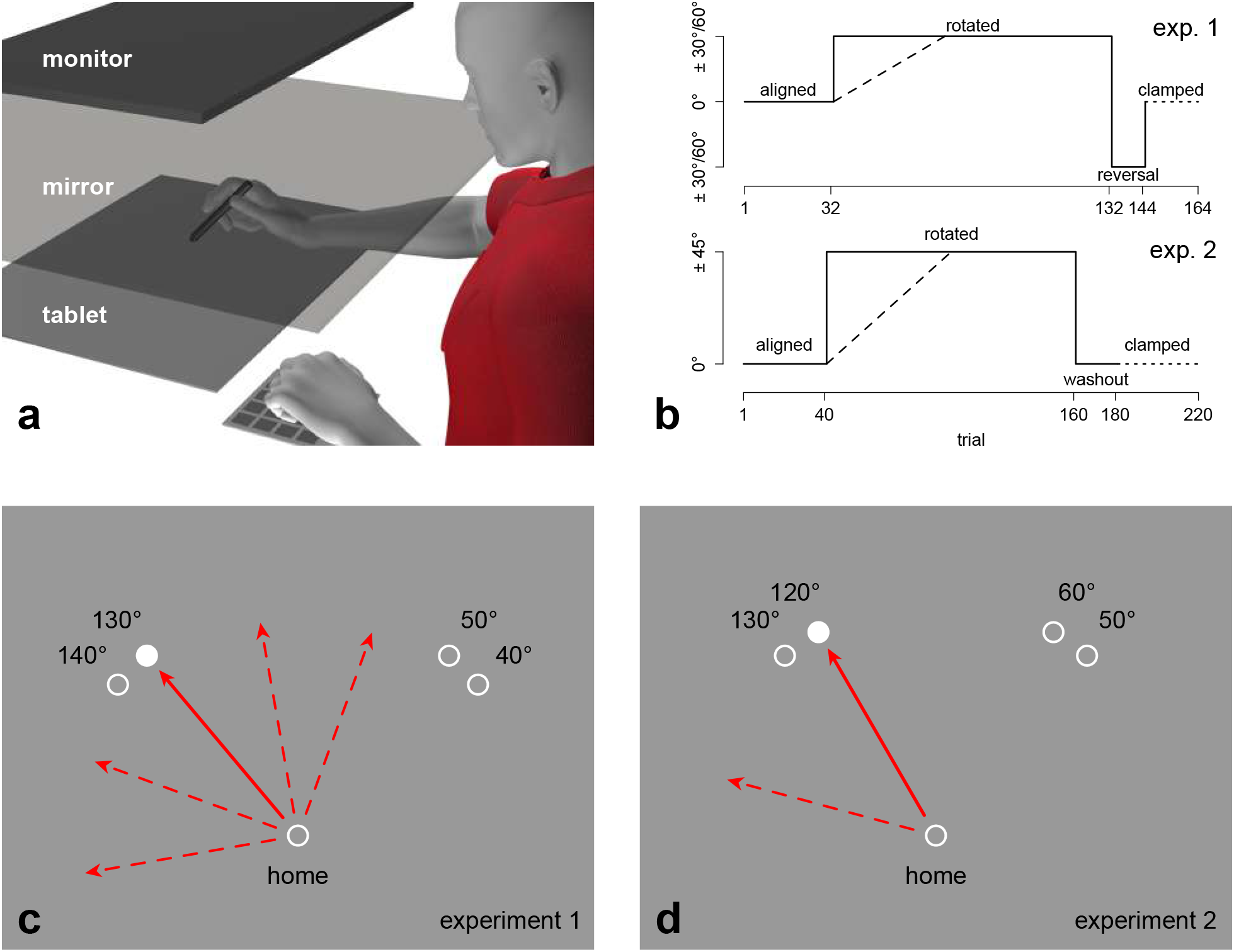
**(a)** Experimental setup. The monitor was located 28 cm above the reflective surface, and the reflective surface was located 26 cm above the tablet. **(b)** Procedure. Overview of the perturbation schedule introduced abruptly and gradually. In experiment 1 (top), a full rotation during the training phase could be either 30° or 60°, and the reversal phase would be of equal magnitude (30° or 60°) and in the opposite direction of the training phase. In experiment 2 (bottom), the training phase is 45°, and the washout phase would have no perturbation. **(c)** Hand-cursor reach from experiment 1. The cursor representing the hand is rotated by 30° or 60°. Visual targets here are presented either 40° or 50° from the midline on either the left or right side of the workspace. The solid red path initially goes straight to the whitetarget, but the cursor veers off by 30° or 60°. Once adapted, this is corrected such that the solid red hand path is moving 30° or 60° in the opposite direction (indicated by the red dashed line) and the cursor is moving straight to the target. **(d)** Hand-cursor reach from experiment 2. Cursor reaches are similar to experiment 1, except the visual targets here are presented either 50° or 60° from the midline, and the cursor is rotated by 45° during training.

#### Procedure

Visuomotor Rotation Task: Participants received continuous visual feedback of their unseen hand position via a white cursor; a 1 cm circle. Participants were instructed to make reaching movements from the home position to the visual target as quickly and accurately as possible. The visual target was a circle with 0.6 cm diameter, and was located 10.4 cm away from the home position. The visual targets were presented either 40° or 50° from the midline on either the left or right side of the workspace (Figure 1c). Once the target was acquired, the trial would end, and the participant would return back to the home position.

Participants performed two visuomotor adaptation tasks sequentially, one where the perturbation during training was introduced abruptly, and one where it was introduced gradually. Both conditions had 4 different phases: aligned, training, reversal, error-clamp (Figure 1b). Both conditions started with the aligned phase, where the cursor represented the true location of the participant’s unseen right hand. During the training phase, the cursor representing the participant’s unseen right hand was rotated around the home position. Participants were asked to make a straight reach to a specific target in the workspace. The cursor representing their actual hand position was then rotated either clockwise or counterclockwise, for which the participant had to reach in the opposite direction to compensate for this perturbation. For the abrupt condition, the cursor was rotated by 30° or 60° for the entire training phase (different groups adapted to the smaller 30° rotation or the larger 60° one). For the gradual condition, the perturbation ramped up to a 30° rotation in increments of 0.75° or to a 60° rotation in increments of 1.5°, such that it took 40 trials to get to the full rotation (in both cases: 2.5% of the rotation per trial), and continued at the full rotation for the remaining 60 trials of the training phase (Figure 1b). During the reversal phase, participants were exposed to an equal rotation in the opposite direction from the training phase. During the final error-clamp phase, the cursor would always move in the direction of the target irrespective of the participant’s actual reach direction. The movement of the cursor along this straight trajectory was still under control of the participant insofar as the distance of the cursor from the home position was kept the same as the distance between the actual hand and the home position. During this phase, participants received no visual feedback of their hand location on the way back to the home position. However, to help participants return to the home position, an arrow at the home position indicating the direction of their actual hand location, guided most of the return to the home position. Once they were near the home position, the cursor representing their unseen hand location would also appear again. In both abrupt and gradual conditions, participants were given 32 trials of an aligned phase, 100 trials of the training phase, 12 trials of a reversal phase, and 20 trials with error-clamped feedback (Figure 1b).

#### Design

For both groups (younger 30°, younger 60°), the experiment was a within-subjects design, such that all participants completed both abrupt and ramped conditions. The experiment began with a familiarization phase, which comprised of 8 aligned trials and 8 error-clamp trials. Next, participants completed one of the two visuomotor adaptation tasks (e.g. abrupt), followed by a mandatory break, and finished by completing the other visuomotor adaptation task (e.g. ramp). The following variables were counterbalanced across participants: the order of the conditions (abrupt or ramp), the side of the workspace that the targets appeared (left or right), and the direction of the rotation (clockwise or counterclockwise). Therefore, participants received one of eight possible variations of the experiment. Counterbalancing the side of the workspace and direction of the rotation was performed to avoid transfer.

## Data Analysis

In order to assess performance throughout the task, for each reaching movement, we calculated the angular reach deviation at the point of maximum hand velocity. Angular reach deviation is the angular difference between a straight line from the start position to the target position, and a straight line intersecting the start position and the position of the participant’s hand. For comparisons that include both 30° and 60° groups, we scaled the angular reach deviation to a proportion of adaptation by dividing it by the rotation size.

### Order Effects

Before addressing the main objectives, we first checked for any order effects from the within-subjects design. We fit a simple asymptotic decay function to each individual participant’s data from the first rotation of the abrupt condition:

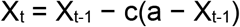

This fits a curve to the motor output (X) with two parameters: an asymptotic level of adaptation (a) and a rate of change (c) in the range [0,1], which expresses how quickly the asymptotic level of adaptation is achieved. Since the rates of change can only be evaluated from the abrupt conditions, we compared rates of change for participants who did this condition first, to those who did it second (after the ramped condition). An ANOVA was performed with the rate of change as the dependent variable, and group (younger 30°, younger 60°) and first condition (abrupt, ramp) as between-subjects factors.

### Asymptotic Level of Adaptation and Rebounds

To assess any reach adaptation differences between the abrupt and ramped conditions, we test reach deviations at two points in the task. First at the end of the first rotated phase, assessing the asymptotic level of learning, which we hypothesize to not be affected as people will have sufficient time to adapt, regardless of how the rotation is introduced. Second, at the end of the zero error-clamped phase. We hypothesize that the reach deviations at this stage are a measure of residual (stable) implicit adaptation, that is decreased by the reversed phase. Hence any differences between conditions should indicate that implicit adaptation was either larger to begin with, or more robust against the effects of the reversal phase in one of the conditions. To test this, we first performed a repeated measures ANOVA on scaled reach deviations, averaged across the last 10 trials of the first rotated phase, and the last 10 trials of the clamped phase. We used block (rotated or clamp) and condition (abrupt or ramped) as within-subject factors and rotation size (30° or 60°) as between-subjects factor. Since we found no significant differences, we also test for equivalence of these conditions by calculating a Bayes Factor (BF_10_) for each block, as the posterior probability of the alternative hypothesis (there is a difference) divided by the posterior probability of the null hypothesis (there is equivalence) given a non-informative prior and the data. A BF_10_=1 means both hypotheses are equally likely. Within the interval 1/3 to 3 (either hypothesis is up to 3 times more likely than the other) there is only anecdotal evidence (21, 22). However, a BF->3 or BF_10_<1/3 (or 0.333) is considered moderate evidence in favor of the alternative hypothesis or the null hypothesis, respectively, whereas values of BF_10_>10 or BF_10_<0.1 are considered strong evidence.

#### Model Fits

In the two-rate model (Smith et al., 2006), the motor output X on trial t (X_t_) is simply the sum of the output of the slow and fast processes on the same trial. Both processes learn from the error on the previous trial (e_t-1_), and retain part of the previous adaptation (X_s,t-1_, X_f,t-1_). The four crucial parameters that are being tested are the learning rates (Ls, Lf), and the retention rates (Rs, Rf) for each of the two processes. The two-rate model integrates the learning and retention rates for each process as follows:

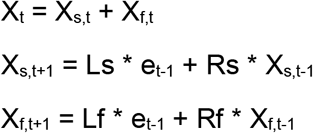

The values for all four parameters (Ls, Lf, Rs, Rf) should fall in the range [0, 1]. To ensure that the fast process learns quicker than the slow process we add the constraint that Lf>Ls, and to ensure the slow process retains more than the fast process, we add the constraint that Rf<Rs.

The model was fitted to the behavioural data of every participant, the group mean and group means across bootstrapped sampling of participants within a group, using a mean square error (MSE) minimization method. To find optimal starting positions for the fitting algorithm, a grid search will be performed on 14^4 points in parameter space, reduced to the parameter combinations in the grid complying with the constraints. We obtain MSE’s for all these parameter combinations, and use the 10 parameter combinations with the lowest MSEs as starting points for a least mean squared error fitting algorithm. The four parameters (Ls, Lf, Rs, Rf) from the fit with the lowest mean squared error were used.

To assess the magnitude of inaccuracies in the model fits, we performed a parameter recovery simulation on the data from younger adults in experiment 2. In this procedure, we use the MSE between the group average data and two-rate model fit as the standard deviation of normally distributed noise (mean 0) applied to the best fitting model and recover the model parameters 1k times, each with different random noise. The obtained 95% confidence intervals for the 8 parameters serve as a range of values where differences between two values of the same parameter are indistinguishable from noise.

The differences between the model parameters of the different conditions were assessed by means of bootstrapping. We make use of the within-subject design by fitting the model to the data of the same samples of participants in each condition, and hence get differences between parameter values from the same random sample of participants. If the 95% confidence intervals for the differences in parameter values overlap with the 95% confidence interval from the parameter recovery simulation, we will treat them as not statistically different.

Experiment 1 was run in Python 2.7 using PyVMEC (https://github.com/thartbm/PyVMEC), and experiment 2 was run in custom Python (participants in Toronto) and Matlab (participants in Munich) code. All data was processed and analyzed using R version 4.0.5. All data can be found on the OSF repository: https://doi.org/10.17605/OSF.IO/CZ3BD. All data processing and analysis can be found at https://github.com/thartbm/GradualTwoRate.

### Experiment 2: Mild Cerebellar Ataxia and Age

#### Participants

Seventy-seven subjects participated in this experiment (thirty younger adults, twenty-five older adults, twenty-two mild cerebellar ataxic patients). The younger adult group consisted of subjects ages 20 +/-2 years old (1 lefthanded, 3 males), the older adult group were ages 60 +/9 years old (1 lefthanded, 14 males), and the mild cerebellar ataxic patients were ages 61 +/-15 years old (all right handed, 9 males). Participants were tested partly at the Centre for Vision Research, York University, Toronto, Canada, and partly at the University Hospital LMU, Munich, Germany. To minimize the effects of extra-cerebellar impairments, recruitment was focused primarily on cerebellar infarcts. Only 7 of the 22 patients (Table 1) had degenerative diseases most likely affecting extracerebellar regions. The experimental setup, the protocol, and the task were identical at both locations. The used hardware differed in only minor details. All participants reported having normal or corrected-to-normal vision. All participants gave prior informed written consent, and were naive to the purpose of the study.

**Table 1:** Mild Cerebellar Ataxic Patients. Information on the years since diagnosis, diagnosis, SARA score, and side of lesion. SARA is the scale of the assessment and rating of ataxia, an 8-item performance based scale resulting in a score between 0 and 40, where 0 is almost no ataxia and 40 is severe ataxia.

| Years Since Diagnosis | Diagnosis | SARA | Laterality |
| --- | --- | --- | --- |
| 34 | infarct superior cerebellar artery | 0.5 | Left-sided lesion |
| 14 | infarct superior cerebellar artery | 3.5 | Left-sided lesion |
| 0 | infarct superior cerebellar artery | 8 | Left-sided lesion |
| 9 | infarct posterior inferior cerebellar artery | 0 | Right-sided lesions |
| 10 | infarct posterior inferior cerebellar artery | 0 | Left-sided lesion |
| 14 | infarct posterior inferior cerebellar artery | 2.5 | Left-sided lesion |
| 11 | infarct posterior inferior cerebellar artery | 3 | Bilateral lesions |
| 0 | infarct posterior inferior cerebellar artery | 5.5 | Left-sided lesion |
| 14 | not further specified cerebellar infarct | 0 | Left-sided lesion |
| 9 | not further specified cerebellar infarct | 0 | Right-sided lesions |
| 12 | not further specified cerebellar infarct | 3.5 | Right-sided lesions |
| 14 | not further specified cerebellar infarct | 4 | Right-sided lesions |
| 0 | not further specified cerebellar infarct | 5 | Left-sided lesion |
| 14 | not further specified cerebellar infarct | 7 | Left-sided lesion |
| 12 | not further specified cerebellar infarct | 9 | Right-sided lesions |
| 37 | cerebellar atrophy | 6.5 | Bilateral lesions |
| 0 | cerebellar atrophy | 19.5 | Left-sided lesion |
| 5 | autosomal-dominant cerebellar ataxia | 8 | Bilateral lesions |
| 16 | autosomal-dominant cerebellar ataxia | 10 | Bilateral lesions |
| 17 | spino-cerebellar ataxia | 5 | Bilateral lesions |
| 16 | episodic cerebellar ataxia type II | 8 | Bilateral lesions |
| 49 | autosomal-dominant cerebellar ataxia | 11 | Bilateral lesions |

#### Apparatus

The equipment used in this experiment was the same as in experiment 1 for the participants in Toronto. For the participants tested in Munich, we used a comparable computer monitor (HPL2245wg, 22”, 60 Hz), mirror, tablet (WACOM Cintiq 21UX, 432 mm × 324 mm), and stylus. Similar to experiment 1, the visual stimuli were projected from the downward facing monitor onto the mirror, such that the stimuli were perceived to be in the same horizontal plane as the tablet below.

#### Procedure

The visuomotor rotation task was similar to Experiment 1, with the main differences being in the perturbation schedule. There were minor differences in the reach amplitude and the target locations were slightly closer to the midline. In this experiment, the rotation size of the perturbation was 45°, each abrupt and gradual condition consisted of 220 trials, and instead of a reversal phase, there was a washout phase. A washout phase is similar to the aligned phase, where there is no rotation of the cursor. This was used to make the experiment doable for patients who would otherwise have had to adapt to a 90° change in rotation between the first and second rotated phases of the task. Both abrupt and gradual conditions had an aligned phase of 40 trials, a training phase of 120 trials, a washout phase of 20 trials, and an error-clamp phase of 40 trials. The training phase of the gradual condition ramped up in increments of 0.75° (or 1⅔% of the rotation per trial), such that it took 60 trials to get to the full 45° rotation, and then continued at the full rotation for the last 60 trials of the phase. The target was located 12 cm away, and presented either 25° or 35° from the midline on either the left or right side of the workspace.

#### Design

This experiment was a within-subjects design, such that all participants completed both abrupt and gradual conditions. After the familiarization phase, participants completed 80 baseline trials: 40 aligned trials and 40 error-clamp trials (10 trials to each target per phase). Next, participants completed one of the two visuomotor adaptation tasks (e.g. abrupt), followed by a mandatory break, and finished by completing the other visuomotor adaptation task (e.g. gradual). The order of the conditions (abrupt or gradual), and the side of the workspace that the targets appeared (left or right) were counterbalanced. The two targets on the left side were presented during tasks with a clockwise rotation, and the two on the right were presented during a counterclockwise rotation. Therefore, participants received one of four possible variations of the experiment.

#### Data Analysis

Same as Experiment 1.

## RESULTS

## Experiment 1: Rotation Size

### Order Effects

Before we could address our main question of whether there were learning differences between the conditions, we first had to rule out any effects of order with our within-subjects design. We found no effect of order on rate of change across the two rotation sizes (F(1,41)=0.496, p=0.523). Hence, we will use all data as is for further analyses.

### Asymptotic Level of Adaptation and Rebounds

Once the order effects had been ruled out, we proceeded to check the asymptotic level of adaptation, and the absolute size of the rebounds in the error-clamp phases. As expected, there was an effect of block (rotated or clamped) showing larger deviations at the end of the rotated block compared to the rebound at the end of clamped block (Figure 2) (F(1,43)=810.707, p<0.001). There was also an effect of rotation size (F(1,43)=18.125, p<0.001) that interacts with block (F(1,43)=13.781, p<0.001) due to a larger rebound in the younger 30° group. However, the main take-away was that there was no main effect of condition (F(1,43)=0.00, p=0.999), nor does condition interact with any of the other variables. This could mean that the way the rotation was introduced (abrupt or ramped) had no effect on the asymptotic level of adaptation here, nor did it affect the size of the spontaneous rebound. A Bayesian analysis provided moderate support for this null hypothesis (BF_10_=0.18).

**Figure 2.**
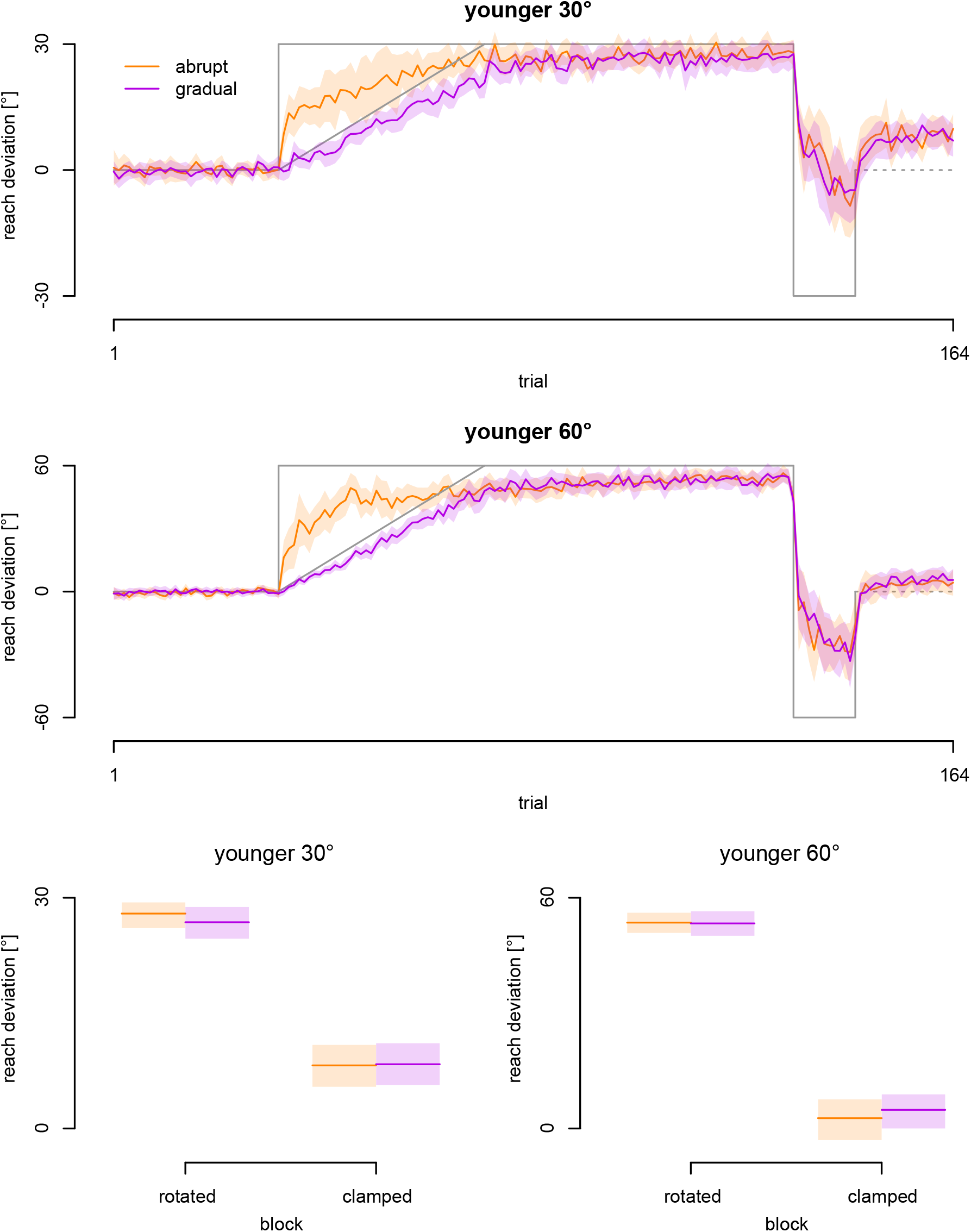
Reach deviations for each of the abrupt and ramped conditions. Top two rows: The data in orange here represents the mean angular reach deviations of the abrupt condition, whereas the data in purple represents the mean angular reach deviations of the ramped condition for groups adapting to the 30° rotation (top) and 60° rotation (middle row). The lightly shaded area in each graph represents the 95% confidence interval. The bottom graphs are the mean reach deviations for the last 10 trials of the rotated and error-clamped blocks for the 30° rotation group on the left and 60° rotation group on the right.

### Two-Rate Model Fits

Lastly, we wanted to see if the two-rate model fits (Figure 3) would be different between the abrupt and ramped conditions. Looking at Figure 4, the distributions of parameter values clearly overlap. The confidence intervals for the difference between the parameter values obtained from the two conditions for both groups, and found that zero was included in all of the confidence intervals. This suggests that the parameters for the two conditions were not significantly different.

**Figure 3.**
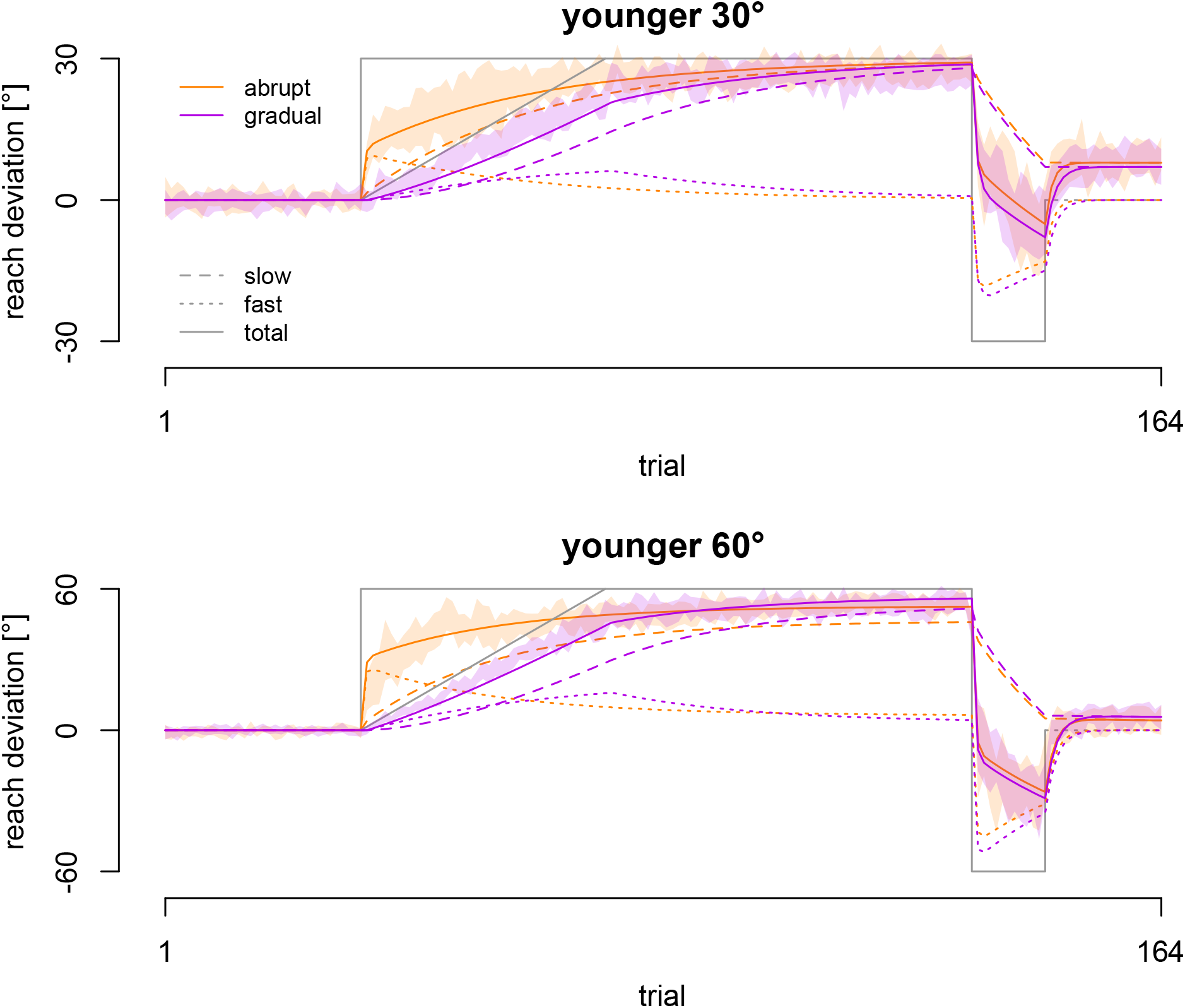
Two-Rate Model Fits. The solid lines here represents the model’s prediction of angular reach deviations: the abrupt condition in orange and the ramped condition in purple. The lightly shaded area in each graph represents the 95% confidence intervals of the reach data, illustrating the goodness of fit. The dashed lines represent the slow processes, whereas the dotted lines represent the fast processes. This data was fit for groups adapting to the 30° rotation (top) and 60° rotation (bottom row).

**Figure 4.**
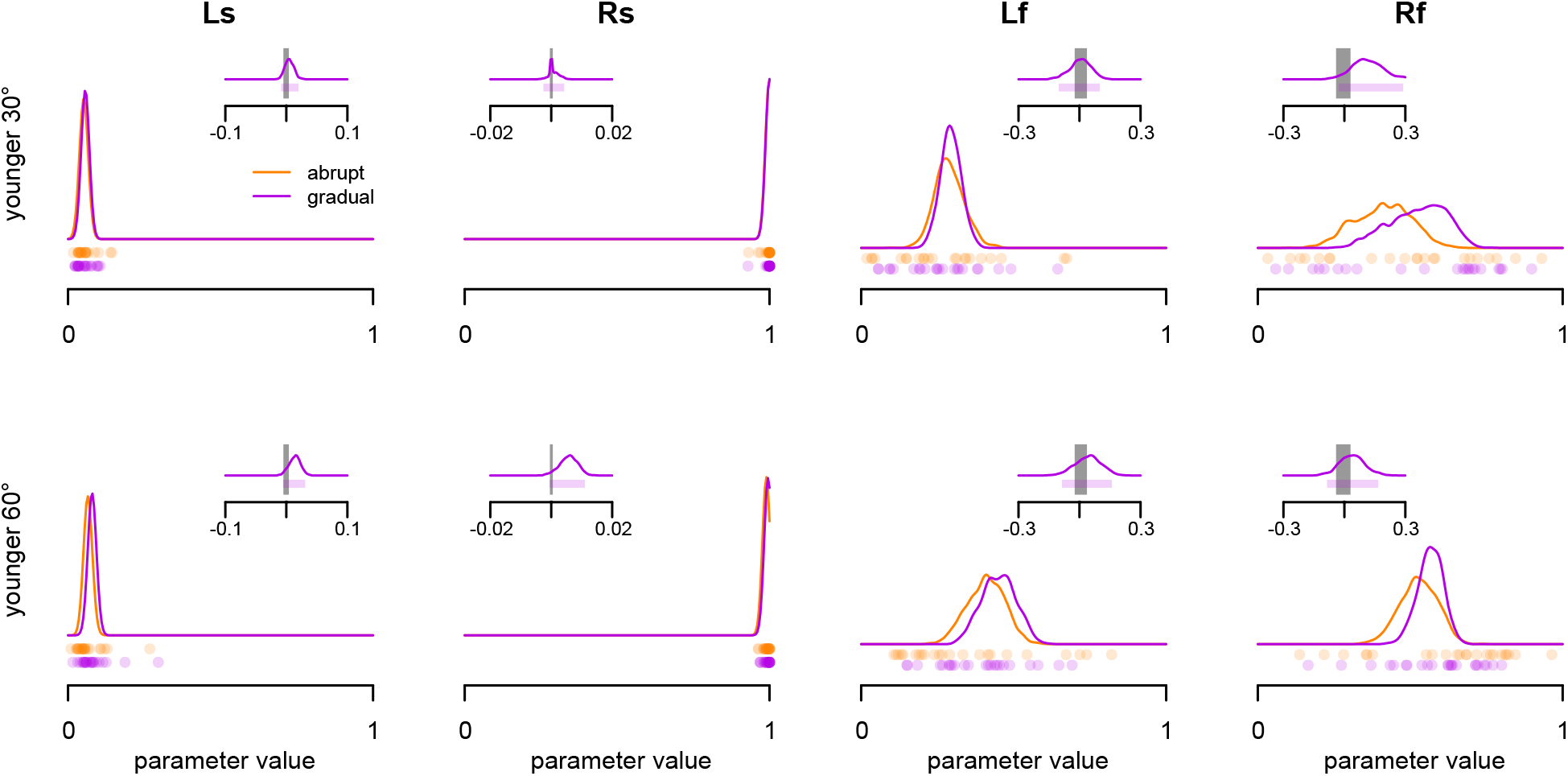
Two-Rate Model Parameter Differences. The orange and purple lines represent the distribution of bootstrapped parameters and the dots are the parameter values for individual participants. The orange is the abrupt condition, and the purple is the ramped. In the small inset, we see a distribution of differences (purple density curve) and 95% confidence interval (purple shaded area). This can be compared to the 95% confidence interval of recovered parameters (gray shaded area) denoting differences that are indistinguishable from noise.

### Experiment 2: Mild Cerebellar Ataxia and Age

#### Order Effects

As before, we first test for an effect of order, by comparing the rate of change in the abrupt condition, as assessed by an exponential decay function with an asymptote, between participants who did the abrupt condition first and who did it second. There was no effect of rotation size (F(2,64)=1.252, p=0.292), no effect of order (F(1,64)=0.763, p=0.385), and no interaction between the two (F(2,64)=0.923, p=0.402). This means there are no order effects, and we will use all data as is.

#### Asymptotic Level of Adaptation and Rebounds

Similarly to Experiment 1, we test the effect of condition (abrupt vs. gradual) on the asymptotic levels of adaptation, and the absolute size of the rebound (Figure 5, bottom row). There was no effect of group (F(2,67)=0.069, p=0.933), or condition (F(1,67)=1.222, p=0.273). As expected, there was an effect of block (F(1,67)=1902.528, p<0.001). Again, condition also did not interact with any of the other factors, and a Bayesian analysis provides moderate support for the null hypothesis that there is no effect of how the rotation is introduced on reach deviations (BF_10_=0.14).

**Figure 5.**
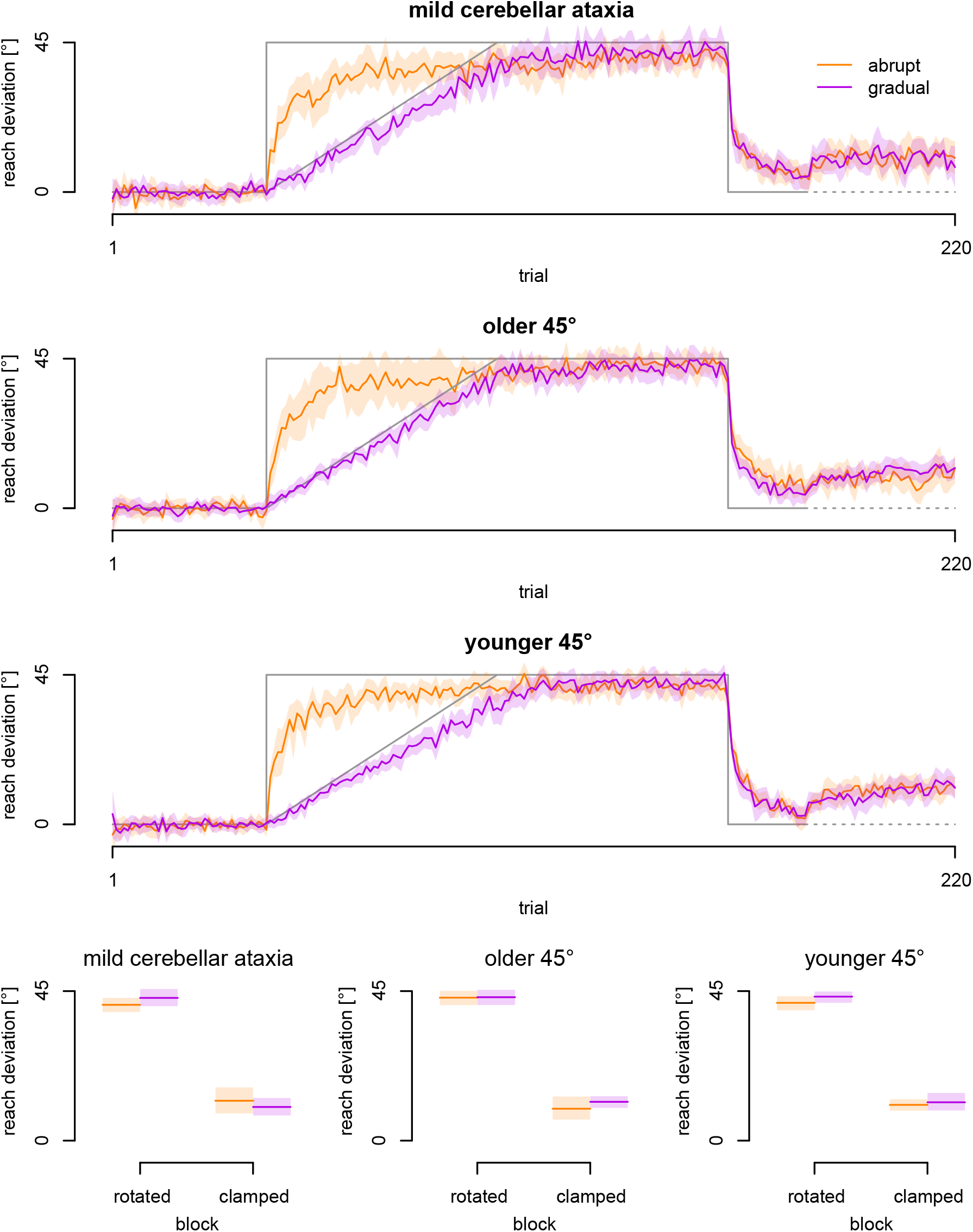
Reach deviations for each of the abrupt and ramped conditions. Top three rows: The data in orange here represents the mean angular reach deviations of the abrupt condition, whereas the data in purple represents the mean angular reach deviations of the ramped condition for the groups adapting to the 45° rotation in the cerebellar ataxic population (top row), the older adult population (second row), and younger adult population (third row). The lightly shaded area in each graph represents the 95% confidence intervals. The bottom graphs are the mean reach deviations for the last 10 trials of the rotated and error-clamped blocks.

**Figure 6.**
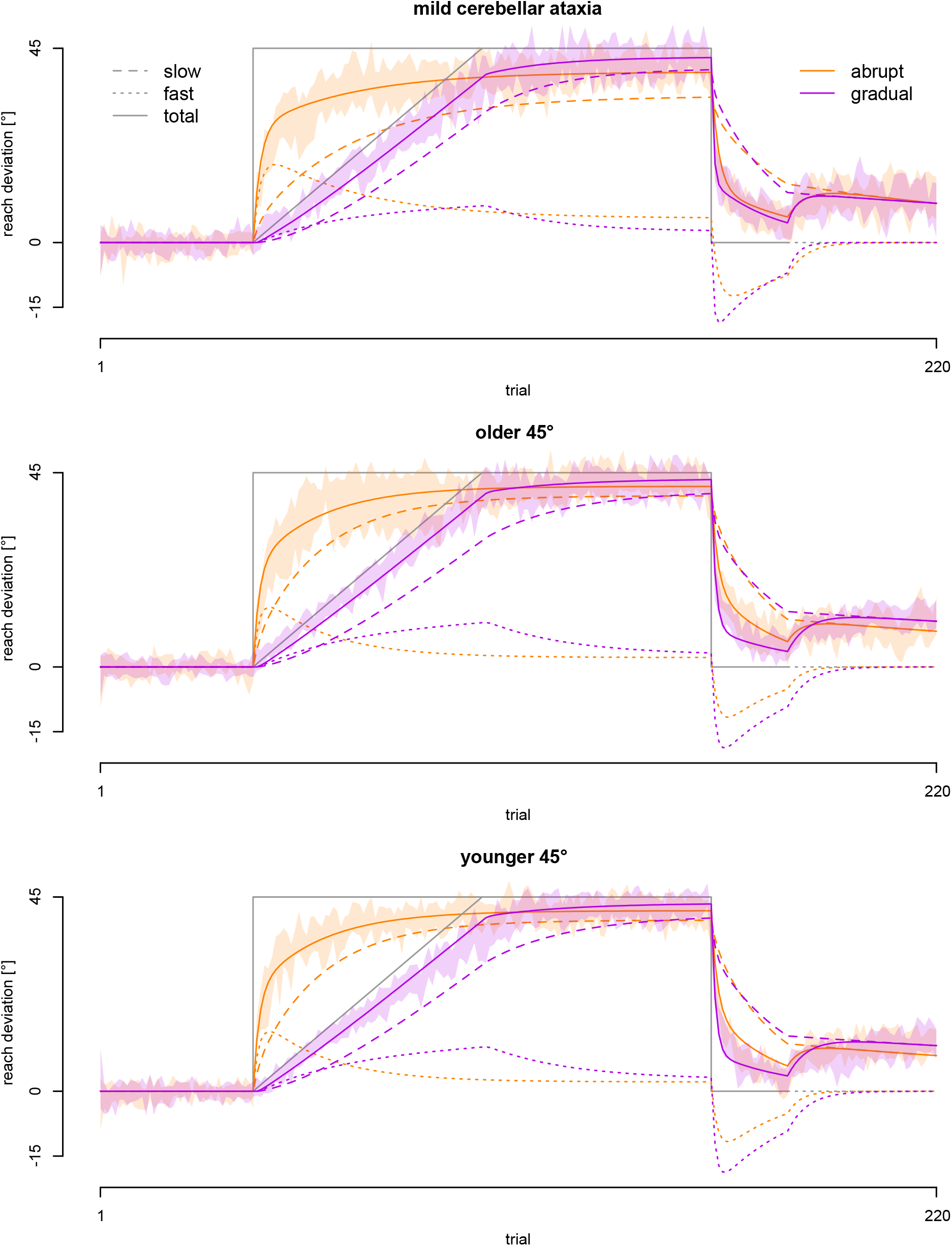
Two-Rate Model Fits. The solid lines here represents the two-rate model’s prediction of angular reach deviations, with the abrupt condition in orange and the ramped condition in purple. The lightly shaded area in each graph represents the 95% confidence intervals of the data, for a visual indication of goodness-of-fit. The dashed lines represent the slow processes, whereas the dotted lines represent the fast processes. This data was fit for the groups adapting to the 45° rotation in the cerebellar ataxic population (top row), the older adult population (middle row), and younger adult population (last row).

#### Two-Rate Model Fits

We looked at the confidence intervals for each of the parameter differences for all the groups and found that for the two older groups (mild cerebellar ataxia, and their age-matched controls), the 95% confidence interval of differences in values of parameter Lf between abrupt and ramped conditions does not include (values indistinguishable from) zero (Figure 7). This shows that in the ramped condition, these groups have higher learning rates for the fast process, although the slow process shows no differences. In the younger group, the fast learning rate shows a small difference between the abrupt and ramped conditions, but the 95% confidence interval for this difference overlaps with the 95% confidence interval for the parameter from the parameter recovery simulation, so it is likely due to chance.

**Figure 7.**
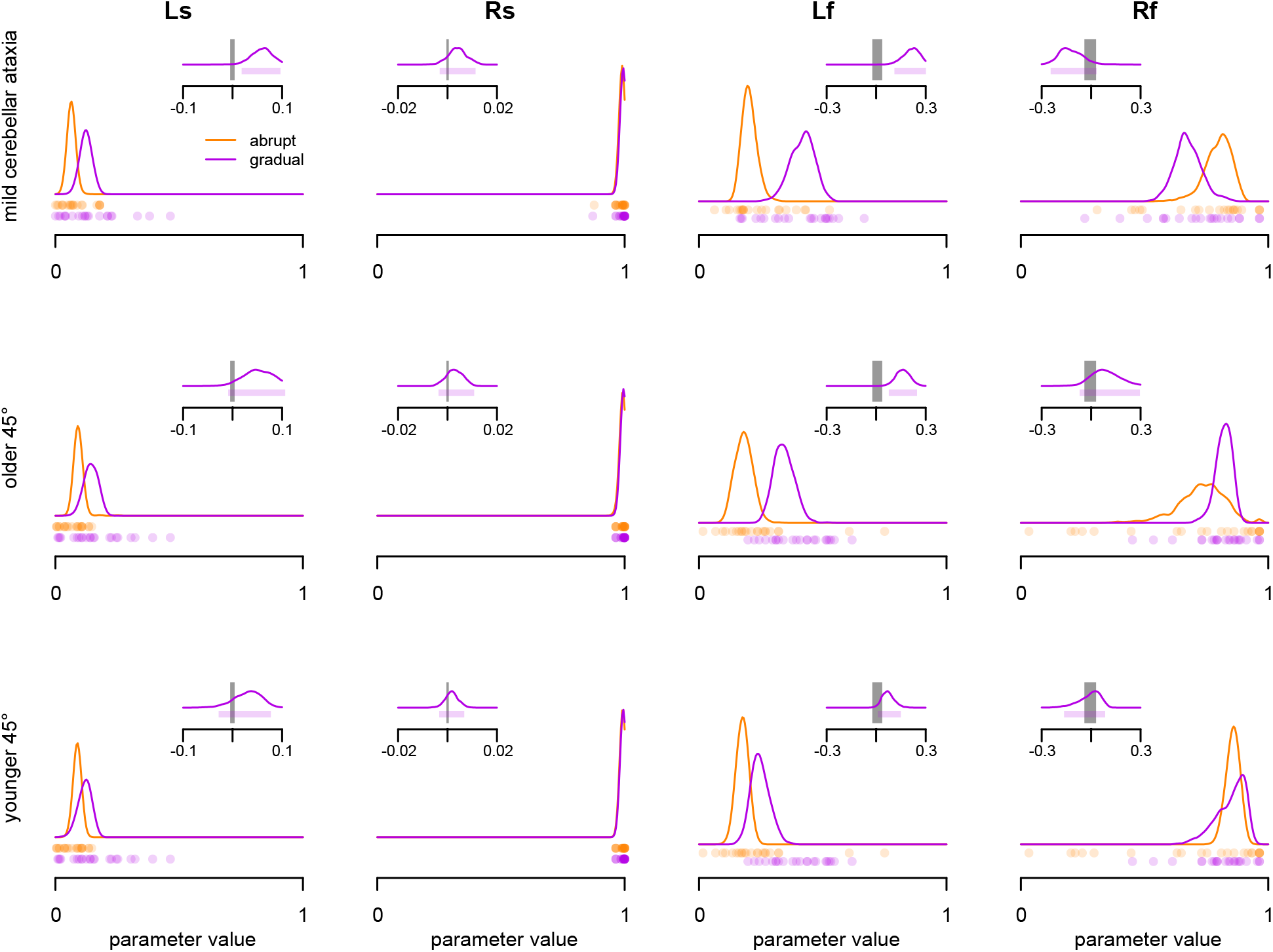
Two-Rate Model Parameter Differences. The lines represent the distribution of bootstrapped parameters and the dots are the parameter values for individual participants. The orange is the abrupt condition, and the purple is the ramped. The insets show distributions of differences as a purple density curve, and the corresponding 95% confidence interval as a shaded purple area. Again, the gray bars are the 95% confidence intervals of parameter differences from the parameter recovery simulation denoting differences that are indistinguishable from noise.

## DISCUSSION

We investigated the reach-adaptation differences when compensating for a perturbation that is introduced abruptly compared to gradually in a visuomotor adaptation. We found that, contrary to our original hypotheses, there were no significant differences in motor adaptation between the abrupt and gradual conditions. This was true in both experiments: (1) when adapting to a 30° and 60° perturbation for younger adults, as well as (2) when adapting to a 45° rotation for younger adults, older adults, and for people with mild cerebellar ataxia. As we will discuss below, although these main findings were not what we were expecting, they do align with previous research and have implications in understanding the processes involved in abrupt versus gradual motor learning.

### Abrupt and Gradual Motor Learning

We found that the way a perturbation was introduced, either abruptly or gradually, had no effect on the rebound or the extent of learning. This was true for all the 30°, 45°, and 60° rotation sizes. The implicit component of motor learning, that is reach aftereffects, is thought to cap at ∼15°, regardless of the rotation magnitude (Kim et al., 2018, Morehead et al., 2017, Modchalingam et al., 2019). The fact that our results show no significant differences in the rebound, as a measure of residual implicit adaptation, between rotation sizes, suggests that the size of the rotation likely does not affect the extent of implicit learning. Given there is no effect of rotation size on implicit learning, a larger rotation likely just recruits more explicit learning (Bond & Taylor, 2015; Heuer & Hegele, 2008; Neville & Cressman, 2018; Werner et al., 2015). This understanding could have implications on how we investigate the explicit and implicit components of learning, and perhaps fast and slow processes of the two-rate model, in the future.

In reviewing the literature, it is evident that the differentiation between these abrupt and gradual conditions, at least behaviourally, is still unclear. Previous research has shown both behavioural (Kagerer et al., 1997, Ingram et al., 2000, Michel et al., 2007, Kluzik et al., 2008) and neurophysiological (Robertson & Miall, 1999, Schlerf et al., 2012, Werner et al., 2014) differences between a perturbation that is introduced abruptly compared to gradually. Although our findings opposed our initial thoughts that the way a perturbation was introduced would affect adaptation performance, previous studies have found similar results as well. In addition to our lab, previous research from other labs also provides evidence to support the idea that the rate at which a perturbation is introduced may not affect adaptation. Previous studies found no difference in reach aftereffects between abruptly and gradually introduced rotation in young adults (Buch et al., 2003, Alhussein et al., 2019), or in typically developed children (Kagerer et al., 2006). The findings from these previous studies are commonly using reach aftereffects once the perturbation has been removed as their measure of adaptation, but there is also research that uses retention as their measure of adaptation. Others have also found no difference in retention between a perturbation that was introduced abruptly compared to gradually in reaching tasks such as a visuomotor hand-cursor adaptation (Coltman et al., 2021) or force-field paradigm (Klassen et al., 2005), as well as in a locomotor adaptation task (Hussain & Morton, 2014). In sum, although our results contradicted our initial hypotheses, there are still several previous studies that have similar findings, that there are no significant behavioural differences in adaptation between a perturbation that was introduced abruptly compared to gradually.

### The Fast and Slow Processes

Our original idea that there would be a greater contribution of implicit adaptation that could be reflected in a larger rebound and potentially a different contribution of the slow process, was based on two assumptions. The first assumption was that abrupt and gradually introduced perturbations elicit different amounts of explicit and implicit learning. The second assumption was that the explicit and implicit components of motor learning map onto the fast and slow processes of the two-rate model (McDougle et al., 2015). Since explicit learning likely depends on large salient errors, when errors are small, or gradually introduced, it might not evoke explicit adaptation and thus mainly drive the implicit component. If there was a greater contribution of the implicit component when a perturbation is introduced gradually, then the residual implicit adaptation as expressed in behavior during the zero error-clamp phase could be larger as well. Similarly, there could have been a greater contribution of the slow process as well. However, neither of these effects were observed. Of course, the model parameters obtained from the fit depend on the magnitude of the rebound, so the assumptions and expectations were somewhat related. In any case, increasing the learning rate of the slow process potentially decreases the rebound, so that any changes in implicit adaptation would more likely manifest as changes in the slow process retention rates, or potentially even in parameters of the fast process. Interestingly, we find weak, inconclusive evidence that in older adults and mild cerebellar ataxia the fast learning rate is affected.Nevertheless, we conclude that at least one, if not both, of our assumptions are likely untrue.

### Motor Learning in Older Adults and People with Cerebellar Ataxia

Similar to the rest of the literature, research in aging and in people with cerebellar ataxia has been mixed. There is evidence to support that there are greater aftereffects with using a gradual compared to abrupt perturbation schedule in people with severe cerebellar ataxia (Criscimagna-Hemminger et al. 2010). Originally, the thought was that aging and cerebellar ataxia may have different contributions of implicit and explicit learning, and therefore modulate the fast and slow processes differently as well. However, a recent paper looking at abrupt and gradual motor learning in cerebellar ataxia had contradicting results (Hulst et al., 2021). Many have also found a lack of support for the idea that motor adaptation is modulated differently for abrupt and gradual perturbation schedules in healthy subjects (Eggert et al., 2021), as well as in cerebellar patients (Gibo et al., 2013; Schlerf et al., 2013; Butcher et al., 2017), and here we find the same. The lack of differences between our patients and the elderly controls also indicates that cerebellar lesions, even if they are accompanied by mild but marked signs of cerebellar ataxia, do not necessarily lead to adaptation deficits. This may also be related to the fact that most of our 15 patients with cerebellar infarcts were in a stable condition long after the first diagnosis (11-8 yrs; Table 1) and may have had sufficient time to develop compensatory mechanisms.

### Limitations and Future Studies

Admittedly, there are difficulties with interpreting null results. One way we enhanced the statistical power in this study was using a within-subjects design to increase the number of participants in each condition. Kagerer et al. (1997) had 5 subjects for each abrupt and gradual condition, Buch et al. (2003) had 5 subjects per condition, Klassen et al. (2005) had 8 subjects per condition, and Kagerer et al. (2006) had 10 subjects perform both conditions. Including all groups, our study had a total of 126 subjects perform both abrupt and gradual conditions. Although there are always struggles with understanding null findings, our within-subjects design and large sample size give some additional strength to the interpretation of our results.

In this study, we did not directly test the explicit and implicit components of motor learning. In future studies, it might be beneficial to add a direct test of explicit and implicit learning during these experiments as well (eg. aiming reports or exclude strategy reaches). Getting this information on the relative contribution of explicit learning could add supplementary evidence for the fact that adapting to a gradual perturbation with small errors is indeed eliciting more implicit learning, compared to an abrupt perturbation with large errors, which should have contributions from both implicit and explicit learning - at least early on in learning and depending on the size of the perturbation. An unpublished study from our lab, however, does test include and exclude strategy open-loop reaches after a perturbation has been introduced in three ways. In the study, gradual and abrupt conditions are compared with a step-wise condition, where the perturbation is introduced in a small number of discrete increases, each followed by a period of consolidation (Modchalingam et al., 2022). Like us, they also found no difference between abrupt and ramped conditions, but the third, “step-wise” method of introducing a perturbation did evoke comparatively more implicit adaptation. This both confirms our current findings and shows that the way a perturbation is introduced can affect how implicitly people adapt to it.

## Conclusions

Our main finding here was that there were no significant differences in adaptation between a perturbation that was introduced abruptly or gradually. This was true across rotation sizes (30°, 45°, 60°), as well as between younger adults, older adults, and people with mild cerebellar ataxia. One major take-away from this study is that maybe we should not equate the fast and slow processes of the two-rate model to the explicit and implicit components of motor learning. As a secondary take-away, the lack of difference in the rebound provides further support for the idea that the size of the rotation may not affect the extent of the slow process that could reflect implicit learning. Although it is not fully settled, our study provides a significant contribution to the conversation about whether adaptation processes are modulated by the way a perturbation is introduced.

## Acknowledgements

This work was supported by the Natural Science and Engineering Research Council. DYPH and AB are supported by the Vision Science to Applications program.

## REFERENCES

Alhussein L, Hosseini EA, Nguyen KP, et al (2019) Dissociating effects of error size, training duration, and amount of adaptation on the ability to retain motor memories. Journal of Neurophysiology 122:2027–2042. https://doi.org/10.1152/jn.00387.2018

Bond KM, Taylor JA (2015) Flexible explicit but rigid implicit learning in a visuomotor adaptation task. Journal of Neurophysiology 113:3836–3849. https://doi.org/10.1152/jn.00009.2015

Buch ER, Young S, Contreras-Vidal JL (2003) Visuomotor Adaptation in Normal Aging. Learn Mem 10:55–63. https://doi.org/10.1101/lm.50303

Butcher PA, Ivry RB, Kuo S-H, et al (2017) The cerebellum does more than sensory prediction error-based learning in sensorimotor adaptation tasks. Journal of Neurophysiology 118:1622–1636. https://doi.org/10.1152/jn.00451.2017

Coltman SK, van Beers RJ, Medendorp WP, Gribble PL (2021) Sensitivity to error during visuomotor adaptation is similarly modulated by abrupt, gradual, and random perturbation schedules. Journal of Neurophysiology 126:934–945. https://doi.org/10.1152/jn.00269.2021

Criscimagna-Hemminger SE, Bastian AJ, Shadmehr R (2010) Size of Error Affects Cerebellar Contributions to Motor Learning. Journal of Neurophysiology 103:2275–2284. https://doi.Org/10.1152/jn.00822.2009

Eggert T, Henriques DYP, ‘t Hart BM, Straube A (2021) Modeling inter-trial variability of pointing movements during visuomotor adaptation. Biol Cybern 115:59–86. https://doi.org/10.1007/s00422-021-00858-w

Gibo TL, Criscimagna-Hemminger SE, Okamura AM, Bastian AJ (2013) Cerebellar motor learning: are environment dynamics more important than error size? Journal of Neurophysiology 110:322–333. https://doi.org/10.1152/jn.00745.2012

Heuer H, Hegele M (2008) Adaptation to visuomotor rotations in younger and older adults. Psychology and Aging 23:190–202. https://doi.org/10.1037/0882-7974.23.1.190

Hull C (2020) Prediction signals in the cerebellum: Beyond supervised motor learning. eLife 9:e54073. https://doi.org/10.7554/eLife.54073

Hulst T, Mamlins A, Frens M, et al (2020) Can we improve slow learning in cerebellar patients? BioRxiv. https://doi.org/10.1101/2020.07.03.185959

Hulst T, van der Geest JN, Thürling M, et al (2015) Ageing shows a pattern of cerebellar degeneration analogous, but not equal, to that in patients suffering from cerebellar degenerative disease. Neuroimage 116:196–206. https://doi.org/10.1016/j.neuroimage.2015.03.084

Hussain SJ, Morton SM (2014) Perturbation schedule does not alter retention of a locomotor adaptation across days. Journal of Neurophysiology 111:2414–2422. https://doi.org/10.1152/jn.00570.2013

ingram HA, van Donkelaar P, Cole J, et al (2000) The role of proprioception and attention in a visuomotor adaptation task. Experimental Brain Research 132:114–126. https://doi.org/10.1007/s002219900322

Jeffreys H (1961) Theory of Probability, 3rd edn. Oxford University Press, Oxford

Kagerer FA, Contreras-Vidal JL, Bo J, Clark JE (2006) Abrupt, but not gradual visuomotor distortion facilitates adaptation in children with developmental coordination disorder. Human Movement Science 25:622–633. https://doi.org/10.1016/j.humov.2006.06.003

Kagerer FA, Contreras-Vidal JL, Stelmach GE (1997) Adaptation to gradual as compared with sudden visuo-motor distortions: Exp Brain Res 115:557–561. https://doi.org/10.1007/PL00005727

Kim HE, Morehead JR, Parvin DE, et al (2018) Invariant errors reveal limitations in motor correction rather than constraints on error sensitivity. Commun Biol 1:19. https://doi.org/10.1038/s42003-018-0021-y

Klassen J, Tong C, Flanagan JR (2005) Learning and recall of incremental kinematic and dynamic sensorimotor transformations. Exp Brain Res 164:250–259. https://doi.org/10.1007/s00221-005-2247-4

Kluzik J, Diedrichsen J, Shadmehr R, Bastian AJ (2008) Reach Adaptation: What Determines Whether We Learn an Internal Model of the Tool or Adapt the Model of Our Arm? Journal of Neurophysiology 100:1455–1464. https://doi.org/10.1152/jn.90334.2008

McDougle SD, Bond KM, Taylor JA (2015) Explicit and Implicit Processes Constitute the Fast and Slow Processes of Sensorimotor Learning. Journal of Neuroscience 35:9568–9579. https://doi.org/10.1523/JNEUROSCI.5061-14.2015

Michel C, Pisella L, Prablanc C, et al (2007) Enhancing Visuomotor Adaptation by Reducing Error Signals: Single-step (Aware) versus Multiple-step (Unaware) Exposure to Wedge Prisms. Journal of Cognitive Neuroscience 19:341–350. https://doi.org/10.1162/jocn.2007.19.2.341

Modchalingam S, Ciccone M, D’Amario S, et al (2022) Adapting to visuomotor rotations in stepped increments increases implicit motor learning. bioRxiv. https://doi.org/10.1101/2022.07.04.498746

Modchalingam S, Vachon CM, ‘t Hart BM, Henriques DYP (2019) The effects of awareness of the perturbation during motor adaptation on hand localization. PLoS ONE 14:e0220884. https://doi.org/10.1371/journal.pone.0220884

Morehead JR, Taylor JA, Parvin DE, Ivry RB (2017) Characteristics of Implicit Sensorimotor Adaptation Revealed by Task-irrelevant Clamped Feedback. Journal of Cognitive Neuroscience 29:1061–1074. https://doi.org/10.1162/jocn_a_01108

Neville K-M, Cressman EK (2018) The influence of awareness on explicit and implicit contributions to visuomotor adaptation over time. Exp Brain Res 236:2047–2059. https://doi.org/10.1007/s00221-018-5282-7

Robertson EM, Miall RC (1999) Visuomotor adaptation during inactivation of the dentate nucleus. NeuroReport 10:1029–1034. https://doi.org/10.1097/00001756-199904060-00025

Rouder JN, Speckman PL, Sun D, et al (2009) Bayesian t tests for accepting and rejecting the null hypothesis. Psychonomic Bulletin & Review 16:225–237. https://doi.org/10.3758/PBR.16.2.225

Schlerf JE, Galea JM, Bastian AJ, Celnik PA (2012) Dynamic Modulation of Cerebellar Excitability for Abrupt, But Not Gradual, Visuomotor Adaptation. Journal of Neuroscience 32:11610–11617. https://doi.org/10.1523/JNEUROSCI.1609-12.2012

Schlerf JE, Xu J, Klemfuss NM, et al (2013) Individuals with cerebellar degeneration show similar adaptation deficits with large and small visuomotor errors. Journal of Neurophysiology 109:1164–1173. https://doi.org/10.1152/jn.00654.2011

Smith MA, Ghazizadeh A, Shadmehr R (2006) Interacting Adaptive Processes with Different Timescales Underlie Short-Term Motor Learning. PLoS Biol 4:e179. https://doi.org/10.1371/journal.pbio.0040179

Taylor JA, Klemfuss NM, Ivry RB (2010) An Explicit Strategy Prevails When the Cerebellum Fails to Compute Movement Errors. Cerebellum 9:580–586. https://doi.org/10.1007/s12311-010-0201-x

Taylor JA, Krakauer JW, Ivry RB (2014) Explicit and Implicit Contributions to Learning in a Sensorimotor Adaptation Task. Journal of Neuroscience 34:3023–3032. https://doi.org/10.1523/JNEUROSCI.3619-13.2014

Vachon CM, Modchalingam S, ‘t Hart BM, Henriques DYP (2020) The effect of age on visuomotor learning processes. PLoS ONE 15:e0239032. https://doi.org/10.1371/journal.pone.0239032

Vandevoorde K, Orban de Xivry J-J (2019) Internal model recalibration does not deteriorate with age while motor adaptation does. Neurobiology of Aging 80:138–153. https://doi.org/10.1016/j.neurobiolaging.2019.03.020

Werner S, Schorn CF, Bock O, et al (2014) Neural correlates of adaptation to gradual and to sudden visuomotor distortions in humans. Exp Brain Res 232:1145–1156. https://doi.org/10.1007/s00221-014-3824-1

Werner S, van Aken BC, Hulst T, et al (2015) Awareness of Sensorimotor Adaptation to Visual Rotations of Different Size. PLoS ONE 10:e0123321. https://doi.org/10.1371/journal.pone.0123321

